# scSNViz: Visualization and Analysis of Cell-Specific Expressed SNVs

**DOI:** 10.1101/2024.05.31.596816

**Authors:** Siera Martinez, Tushar Sharma, Luke Johnson, Allen Kim, Vania Ballesteros Prieto, Hovhannes Arestakesyan, Sunisha Harish, Jewel Dias, Joseph Goldfrank, Nathan Edwards, Anelia Horvath

## Abstract

**Summary:** Accurately characterizing expressed genetic variation at the single-cell level is essential for understanding transcriptional heterogeneity, allelic regulation, and mutational dynamics within complex tissues. However, few tools enable comprehensive visualization and quantitative analysis of expressed variants across individual cells.

scSNViz is an R package for the exploration, quantification, and visualization of expressed single-nucleotide variants (SNVs) from cell-barcoded single-cell RNA sequencing (scRNA-seq) data. The software supports estimation of variant allele fractions, clustering of SNV expression profiles, and 2D and 3Dl visualization of individual SNVs or user-defined SNV groups. Beyond visualization, scSNViz facilitates investigation of cell-, cluster-, or lineage-specific variant expression patterns, as well as allelic dynamics including imprinting, random allele inactivation, and transcriptional bursting. It interoperates seamlessly with established single-cell frameworks - Seurat for clustering, Slingshot for trajectory inference, scType for cell-type annotation, and CopyKat for copy-number profiling - enabling integrative multi-omic analyses of expressed variation.

**Availability and Implementation:** scSNViz is implemented in R and freely available at https://github.com/HorvathLab/scSNViz (DOI: 10.5281/zenodo.17307516). The package includes comprehensive documentation and example workflows designed for users with limited bioinformatics experience.

## Introduction

Cell-specific expressed single nucleotide variants (SNVs) include DNA-transcribed germline and somatic variants, as well as context-dependent RNA-originating variants, all of which dynamically contribute to transcriptional heterogeneity. These SNVs can influence protein function, gene expression regulation, transcriptional and translational efficiency, and cellular signaling. Mapping SNV distribution and expression at single-cell resolution offers valuable insights into cellular diversity and regulatory mechanisms.

With the increasing availability of scRNA-seq datasets, growing recognition of SNVs contribution to cellular heterogeneity, and rapid emergence of detection tools (Liu *et al*., 2019; Prashant, Liu, *et al*., 2021; Edwards *et al*., 2023; Zhang *et al*., 2023;Zheng *et al*., 2017; Muyas *et al*., 2024; Singer *et al*., 2018), there is a need for dedicated tools to enable in-depth exploration.

We introduce scSNViz, a dedicated tool for visualization, analysis, and graphical representation of SNV patterns in cell-barcoded scRNA-seq data (e.g., 10x Genomics Zheng et al., 2017). scSNViz offers a core suite of functionalities, including quantitative analysis of SNV expression, 2D and 3D visualization of individual or sets (user-defined groups) of SNVs, SNV clustering based on their expression profiles across cells, and comparative analysis across multiple samples. To facilitate integrated analysis within the broader transcriptomic context, scSNViz interfaces with tools such as Seurat (Butler *et al*., 2018), for cell-level gene-expression processing, Slingshot (Street *et al*., 2018), for trajectory inference, scType (Ianevski *et al*., 2022) for cell type annotation, and CopyKat (Gao et al., 2021) for copy number variation profiling. It offers flexible input options, including SCReadCounts outputs (Prashant, Alomran, *et al*., 2021) and user-defined SNV lists.

We applied scSNViz to explore diverse biological contexts across 28 publicly available primary tumor and normal tissue samples (Supplementary Table 1), including: prostate cancer (pc, Ma et al., 2020), non-small cell lung carcinoma (nsclc, Wang et al., 2019, cholangiocarcinoma (chlg, Zhang et al., 2020, and a combined neuroblastoma (nb), normal fetal adrenal (fa), and normal embryo (ne) cohort, Dong *et al*., 2020).

## Software Description

scSNViz is an open-source R package that enables users to explore SNV-related cellular heterogeneity and integrate SNV information into existing scRNA-seq workflows (Figure 1a).

**Figure 1.**
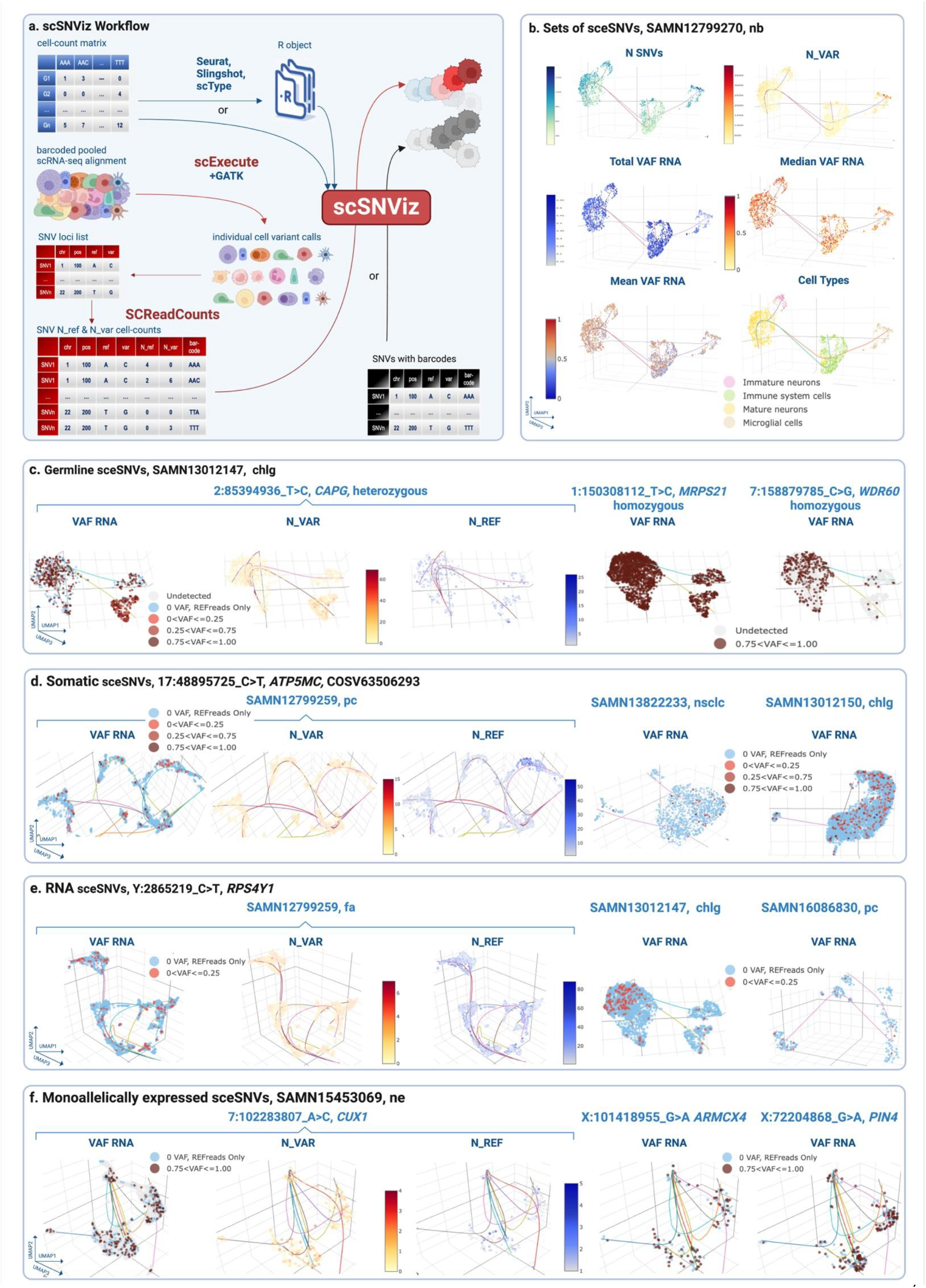
a. scSNViz workflow: scSNViz accepts either raw or processed gene-cell expression values as its first input, and a list of SNVs with cellular barcodes as its second input. It is recommended that the list of SNVs be processed through SCReadCounts, facilitating quantitative visualization based on the number of reference and variant read counts, expressed as color gradients. SCReadCounts-processed data also enables the distinction between SNV loci with solely reference read counts, solely variant read counts, and no read counts, thus allowing for the differentiation between cells with monoallelic reference expression, monoallelic variant expression. The software tools developed by our team and focused on variant analysis from single cells are shown in red. **b. Sets of SNVs:** Overall statistical metrics, accompanied by cell type classification by scType (offered as an option in scSNViz). To enhance visualization, users can select from multiple customizable color designs, including mono- and bi-chromatic gradients. The visualization is exemplified on the publicly accessible sample SAMN12799270, isolated from neuroblastoma tumor tissue; the list of SNVs used is shown in Supplementary Table 2. **c. scSNViz UMAP visualization of individual SNVs with likely germline origin** in sample SAMN13012147 (cholangiocarcinoma primary tumor). For each SNV in the submitted list, VAF_RNA, N_VAR, and N_REF are visualized to evaluate allele-specific expression. Distinct patterns differentiate heterozygous from homozygous SNVs, with homozygous variants showing exclusive expression of the variant allele - enabling clear discrimination based on allelic configuration. **d. scSNViz UMAP visualization of individual SNV 17:48895725 C>T in ATP5MC1, corresponding to the previously reported somatic mutation COSV63506293**, across three samples from different tumors: prostate cancer, non-small cell lung carcinoma and cholangiocarcinoma. In all samples, similar bi-allelic expression patterns were observed. **e. scSNViz UMAP visualization of individual SNV Y:2865219_C>T in RPS4Y1, with likely RNA origin**, across three samples from different tissues: normal fetal adrenal, cholangiocarcinoma and prostate cancer. In all samples, similar expression patterns were observed, with low N_VAR and VAF_RNA values. **f. scSNViz UMAP visualization of individual SNVs from heterozygous loci exhibiting random monoallelic expression at the single-cell level**. These patterns are characterized by a similar number of cells expressing either the variant or the reference allele, typically supported by low read counts (often fewer than 10). Such expression profiles are consistent with both X-chromosome inactivation and transcriptional bursting.

### Input Data Requirements

scSNViz requires two key inputs. A genes-by-cells expression matrix can be provided as a Seurat object - after custom processing such as quality control, scaling, and normalization - or as raw gene expression values from sequencing aligners such as Cell Ranger (Zheng *et al.*, 2017) or STARsolo (Kaminow *et al.*, 2021). In the latter case, scSNViz processes the data using default workflows from Seurat and Slingshot, with optional cell type classification via scType and aneuploidy assessment using CopyKat.

The second input is a tab-delimited list of SNVs with corresponding cellular barcodes. For quantitative visualization, users are encouraged to preprocess this list with SCReadCounts, a tool optimized for barcoded scRNA-seq data that quantifies reference and variant read counts at the single-cell level across all SNVs in a sample. This enables clear distinction between absence of variant expression (negative signal) and absence of gene expression (no signal) in a given cell (Prashant, Alomran, *et al.*, 2021). To ensure compatibility with diverse preprocessing workflows, scSNViz also accepts a simplified input format containing only SNV identifiers, cellular barcodes, and corresponding variant (N_VAR) and reference (N_REF) read counts, facilitating integration with custom pipelines or third-party tools. This format additionally supports categorical representations of SNV presence. scSNViz processes N_VAR and N_REF, calculates the expressed Variant Allele Frequency (VAF_RNA = N_VAR/(N_VAR + N_REF)), and visualizes these three metrics quantitatively by mapping them onto color gradients.

### Global and Cluster-Level Metrics for Sets of SNVs

For sets of SNVs, scSNViz computes and visualizes key metrics that characterize expression patterns across individual cells and clusters. These include N_SNVs (total number of expressed SNVs per cell), N_VAR, N_REF, and total, mean and median VAF_RNA, which are calculated across the entire set of selected SNVs per cell to capture variant expression magnitude and variability (Figure 1b). Total VAF_RNA is computed by summing N_VAR across all selected SNV loci and dividing by the total reads covering these loci (N_VAR + N_REF), providing an overall measure of SNV expression magnitude at the cell-population level. In contrast, median and mean VAF_RNA summarize the central tendency and variability of variant expression across individual cells, offering insights into cell-to-cell heterogeneity. scSNViz also visualizes the distribution of these metrics using histograms, enabling rapid comparison across cell populations or clusters (Supplementary Figure 1).

### Individual SNVs

In addition to global SNV metrics, scSNViz visualizes N_VAR, N_REF, and VAF_RNA for each individual SNV in the submitted set (Figure 1c-f, see below “Examples of Applications”). These metrics provide distinct insights into SNV expression and support the study of various biological phenomena, including global, cell- and cluster-specific expression (Supplementary Figure 2), allele-preferential expression, and gene expression regulation.

### SNV Clustering Analysis

Beyond cell-level clustering, scSNViz also supports clustering of SNVs based on their expression profiles across cells. This is achieved by concatenating N_VAR, N_REF, and VAF_RNA values, followed by transposing the matrix to treat cells as features - forming the “Transposed SNV Matrix.” Dimensionality reduction and clustering are then applied to group SNVs with similar transcriptional activity patterns across the cell population (Supplementary Figure 3). This structure enables identification of SNVs with shared expression profiles across cells, including separating homozygous from heterozygous germline variants.

### Integration of Multiple Samples

**scSNViz** supports SNV visualization across multiple samples using the “enable_integrated” option, which projects SNVs onto a shared dimensionality reduction space via Seurat’s integration method. This approach generates an additional UMAP plot where cells are grouped by sample ID, allowing selective inclusion or exclusion of samples (Supplementary Figure 4). This functionality aids in identifying shared versus sample-specific SNV expression patterns.

### Interactive and Customizable Visualization

scSNViz enables interactive 3D visualization of reduced-dimensionality data, allowing users to dynamically inspect clusters and export 2D snapshots. In addition, scSNViz incorporates user-defined filtering thresholds for expression (e.g., excluding SNVs detected in fewer than 20 cells) and read coverage (e.g., filtering out SNVs with sequencing depth <3 reads). These filters help remove low-confidence SNVs and prioritize biologically meaningful signals. For instance, SNVs with consistently high N_VAR in a specific cluster may reflect a strong cell-type-specific association, while those with low N_VAR may represent technical noise or stochastic transcription. Furthermore, scSNViz offers customizable plotting parameters, including colors, legends, labels, axes, cell sizes, and borders (See Figure 1b). It also provides a clickable option to selectively visualize cells based on VAF_RNA, N_VAR, and N_REF values or specific cell types, with the ability to include or exclude Slingshot trajectories (Supplementary Figure 5). Additionally, scSNViz supports flexible dimensionality reduction methods, including UMAP, t-SNE, and PCA (Supplementary Figure 6).

### Availability and Implementation

scSNViz is distributed as an open-source R package (https://github.com/HorvathLab/scSNViz, DOI: https://doi.org/10.5281/zenodo.17307516) with built-in documentation and an example dataset to illustrate parameter optimization and package use. The toolkit runs efficiently across diverse computing environments, including both high-performance clusters and standard laptops, with typical runtimes in the range of minutes. Runtime scales with the number of cells and features in the input matrix, with the optional use of CopyKat contributing the most overhead - extending runtimes from 5 minutes to several hours for large datasets - though this is readily mitigated by enabling its built-in parallelization, recommended for efficient large-scale analysis (Supplementary Figure 7).

## Examples of Applications

scSNViz can be applied to diverse biological scenarios, providing insights into the functional and regulatory dynamics of SNVs across different cellular contexts, with three examples demonstrated below.

### Inferring Variant Origin from scRNA-seq Data

As most scRNA-seq datasets lack cell-matched DNA data, the probable SNVs origin - germline, somatic, or RNA - can be inferred by analyzing scSNViz-generated N_VAR, N_REF, and VAF_RNA visuals. Germline SNVs, assuming no allele-preferential bias, exhibit mono- or biallelic expression depending on zygosity. Heterozygous variants typically show balanced allele expression, with comparable N_VAR and N_REF values and VAF_RNA centered around 0.5; whereas homozygous variants are expected to display monoallelic expression in all expressing cells (Figure 1c). Somatic SNVs, on the other hand, are often expressed from a single allele in a subset of the cells with a lower, but transcription-consistent VAF_RNA values (Figure 1d). Contrasting both scenarios above, RNA-originating SNVs, such as those resulting from RNA editing (Rees and Liu, 2018) or transcriptional infidelity (Gordon *et al.*, 2009), can display transcription-inconsistent and cell-type-specific patterns, often characterized by low N_VAR and VAF_RNA values (Figure 1e).

### Identifying Novel Patterns in COSMIC reported loci

We applied **scSNViz** to analyze loci previously reported to harbor somatic SNVs in the Catalogue of Somatic Mutations COSMIC (Tate *et al.*, 2019). Notably, several SNVs cataloged as rare or isolated somatic mutations - previously observed in only one or a few patients - were detected across multiple samples in our dataset. For instance, COSV63506293 (17:48895725 C>T in *ATP5MC1*) was identified in 23 out of 28 samples (Figure 1d). Inspection using Integrated Genomic Viewer (IGV, Robinson et al., 2023) confirmed high-quality alignments and high-confidence variant calls for these observations (Supplementary Figure 8).

Interestingly, in most cases this SNV exhibited low N_VAR and low VAF_RNA values. While not excluding a somatic origin, such expression patterns are also consistent with RNA-origin mechanisms, such as RNA editing or transcriptional infidelity. The widespread recurrence of these variants across samples - albeit in a small population of cells - warrants further investigation.

### Monoallelic Expression from heterozygous loci

scSNViz provides a framework for assessing biologically regulated monoallelic expression, including X-chromosome inactivation, where one X chromosome is epigenetically silenced in female cells (Fang *et al*., 2019), and transcriptional bursting, a phenomenon in which genes are transcribed in stochastic pulses (Tunnacliffe and Chubb, 2020). In both scenarios, allele-specific expression at heterozygous loci is expected to yield VAF_RNA values near 0 or 1, reflecting exclusive expression from either the maternal or paternal allele.

Indeed, as expected, we observed widespread monoallelic expression among X-linked SNVs, as well as among numerous heterozygous autosomal SNVs. In many of these cases, both N_VAR and N_REF were low (often fewer than 10 reads) yet showed apparent allelic asymmetry (Figure 1f), consistent with either transcriptional bursting or X-chromosome inactivation. These observations highlight how scSNViz facilitates the visualization of allelic expression imbalance, while emphasizing that low-coverage loci should be interpreted with caution, as technical variability and stochastic transcription can contribute to apparent skew.

Beyond these examples, scSNViz is a versatile tool supporting a wide range of allele-specific investigations, from the dynamics of specific mutations and genes to broader biological processes, including genomic imprinting, clonal lineage tracing, gene expression regulation, and SNV-gene expression relationships.

## Discussion

scSNViz is intended for researchers investigating cell-level DNA and/or RNA-originating nucleotide variance across various biological contexts. The tool offers a user-friendly interface designed for custom visualization of expressed genetic variance derived from scRNA-seq data for both individual SNVs and sets of SNVs. The visualization of both sets and individual SNVs offers distinct information value, providing insights into overall expression patterns and specific allele behaviors, respectively.

We note that SNVs and SNV-sets for analyses may be either called directly or inferred. In our workflow, we perform cell-level variant calling using SCExecute (Edwards et al., 2023) in combination with GATK and Strelka (McKenna et al., 2010; Kim et al., 2018), followed by N_VAR and N_REF assessments using SCReadCounts (see Supplementary Materials and Methods). Variants processed through SCReadCounts can include any genomic positions of interest, such as somatic mutational hotspots or RNA-editing sites, where independent variant calling may not be required.

The selection of SNVs as a set for scSNViz analysis is crucial, as different sets provide distinct insights into transcriptional heterogeneity, including biallelic germline expression, somatic mutation evolution, or RNA-originating variance. For example, germline, somatic, and RNA-originating variants display unique patterns when analyzed as separate sets (Supplementary Figure 9), whereas their combined analysis may obscure these differences.

Given the current understudied nature of SNVs from cell-barcoded scRNA-seq data, we anticipate that scSNViz will drive substantial advancements in data generation, analysis, and interpretation of expressed nucleotide variation. Of note, scSNViz is directly applicable to long read scRNA-seq data, including the emerging datasets produced by Nanopore and PacBio which are expected to significantly expand the number and types of mutations identifiable from scRNA-seq data.

## Authors Contribution

AH developed the concept and drafted the manuscript; SM, TS, AK, and LJ implemented the R package; VBP, SH, JD, JG and NE tested and optimized the software; HA, VBP and JD reanalyzed the data, provided examples of applications, and created additional figures. AH devised and supervised the study. All authors have read and approved the final manuscript.

## Funding

This work has been supported by NCI award R21CA271066 to Anelia Horvath and by McCormick Genomic and Proteomic Center at George Washington University.

## Notes

https://github.com/HorvathLab/scSNViz

### Competing Interest Statement

The authors have declared no competing interest.

### Summary of Updates

ScSNViz has been substantially improved to enhance usability, structure, and functionality. The tool has now been restructured as an R package, increasing modularity and ensuring seamless interoperability with existing R-based workflows. New analytical features have been incorporated, including SNV-level clustering with integrated visual outputs, which are now reflected in both the main manuscript and supplementary materials. Documentation has been significantly expanded, with clearer installation instructions, structured usage examples, and comprehensive guidance suitable for users with varying levels of experience. The manuscript has also been updated to include additional applied examples demonstrating real-world use cases of ScSNViz. Finally, the presentation has been refined: the title has been streamlined, the Software Description section has been reorganized with new subheadings for clarity, an Examples of Applications section has been added, and terminology has been standardized by replacing sceSNV with SNV to avoid confusion with the tool name.

