## Supplementary Figures for "scSNViz: Visualization and Analysis of Cell-Specific Expressed SNVs"

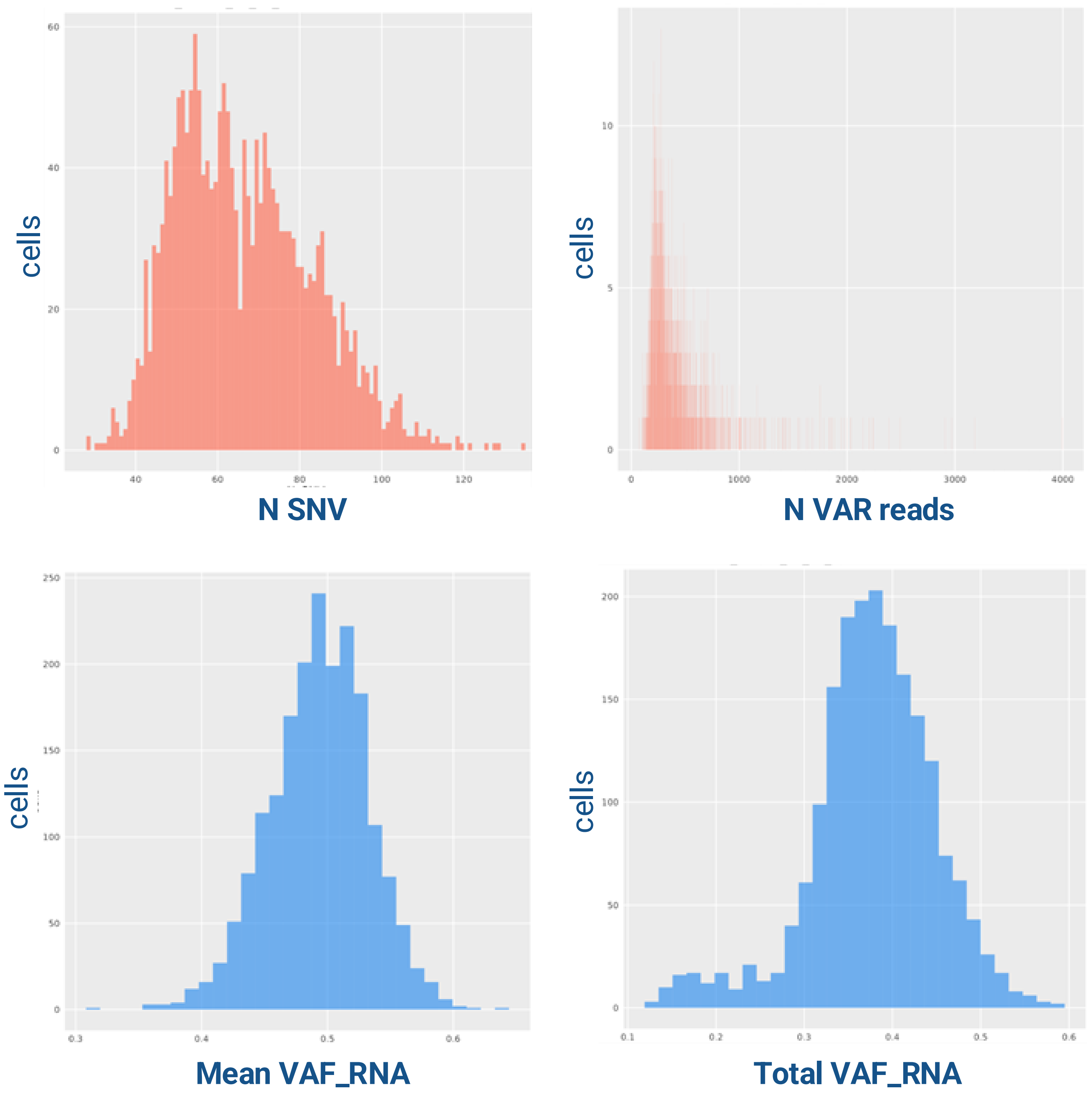


**Supplementary Figure 1**. Histograms depicting metric distribution patterns within bins of cell numbers for the entire set of submitted sceSNVs. This visualization is designed to support a summary-level exploration of cellular heterogeneity.

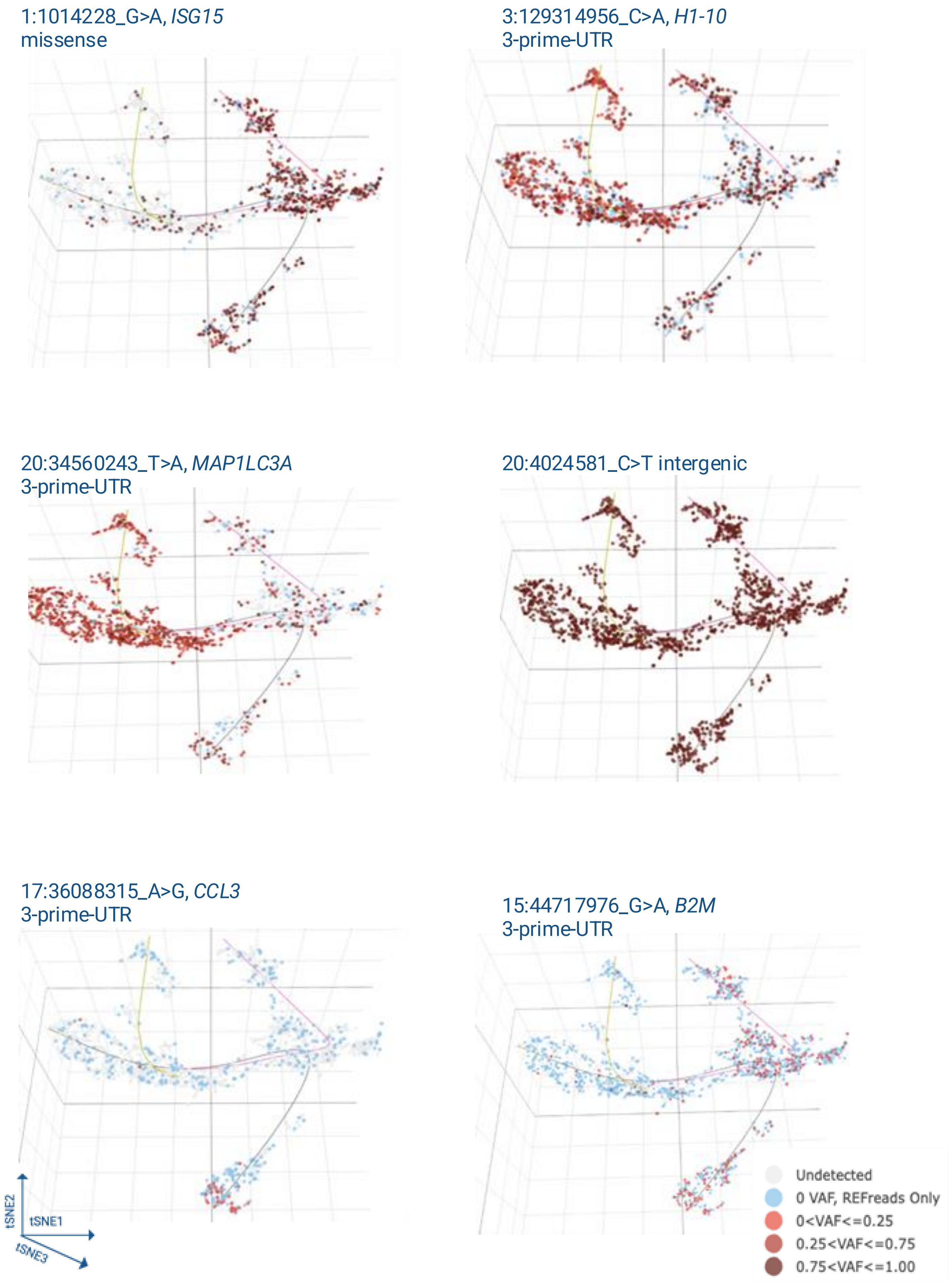


**Supplementary Figure 2**. Examples of different cellular distribution and expression patterns of sceSNV expression. Some sceSNVs show specific expression in certain cell clusters or sub-clusters, while others are expressed across a broader range of cells in the sample, with varying levels of the variant allele in different cell clusters.

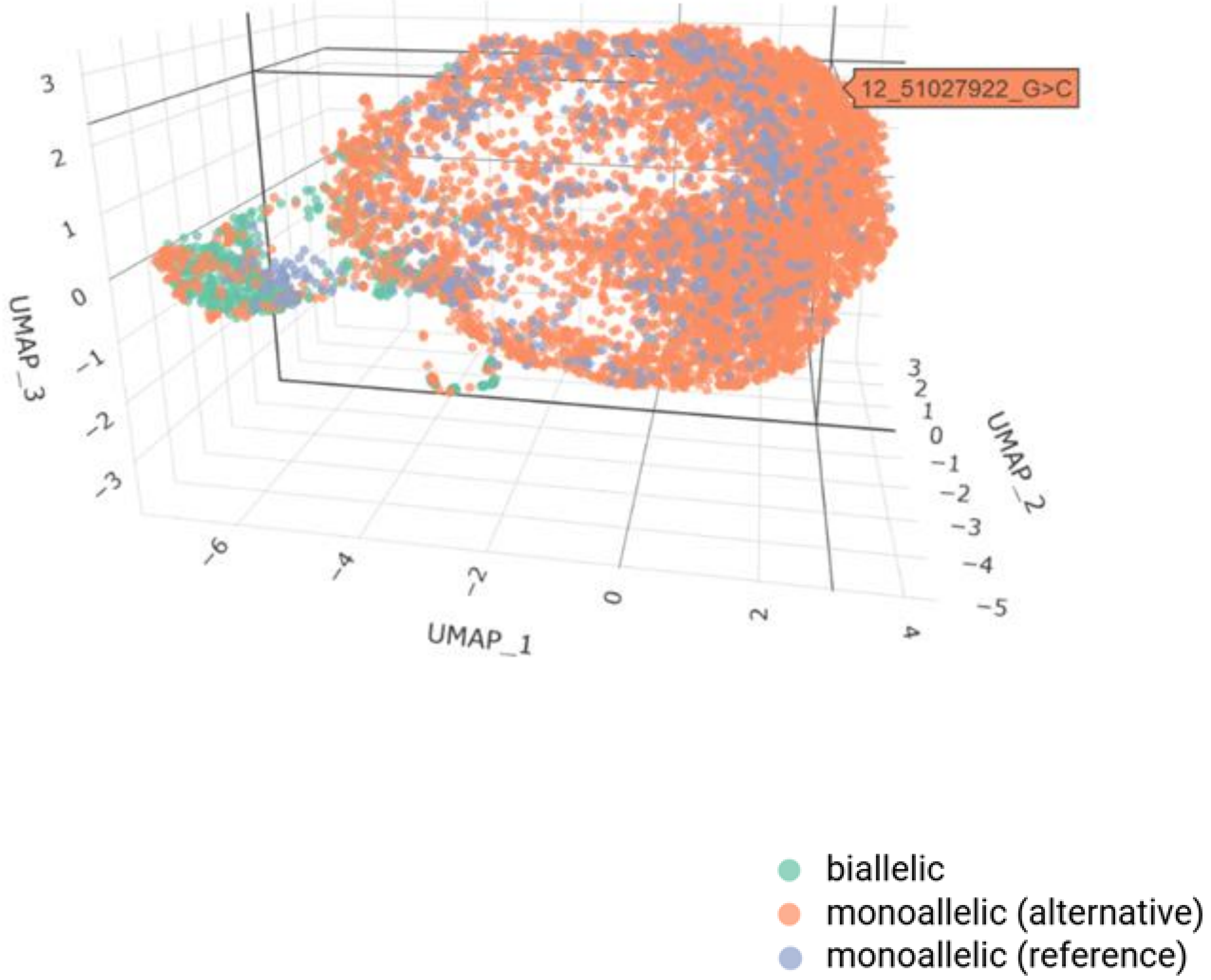


**Supplementary Figure 3**. UMAP visualization of sceSNVs in sample SAMN16086030 based on their expression patterns across cells, generated using the Transpose_SNV_Matrix function in scSNViz. The selected 3D rotation highlights separation between monoallelic sceSNVs (likely homozygous, shown in orange and blue) and biallelic sceSNVs (likely heterozygous, shown in green). Interactive visualization allows users to hover over each sceSNV to display its chromosomal position and variant ID of selected sceSNVs.

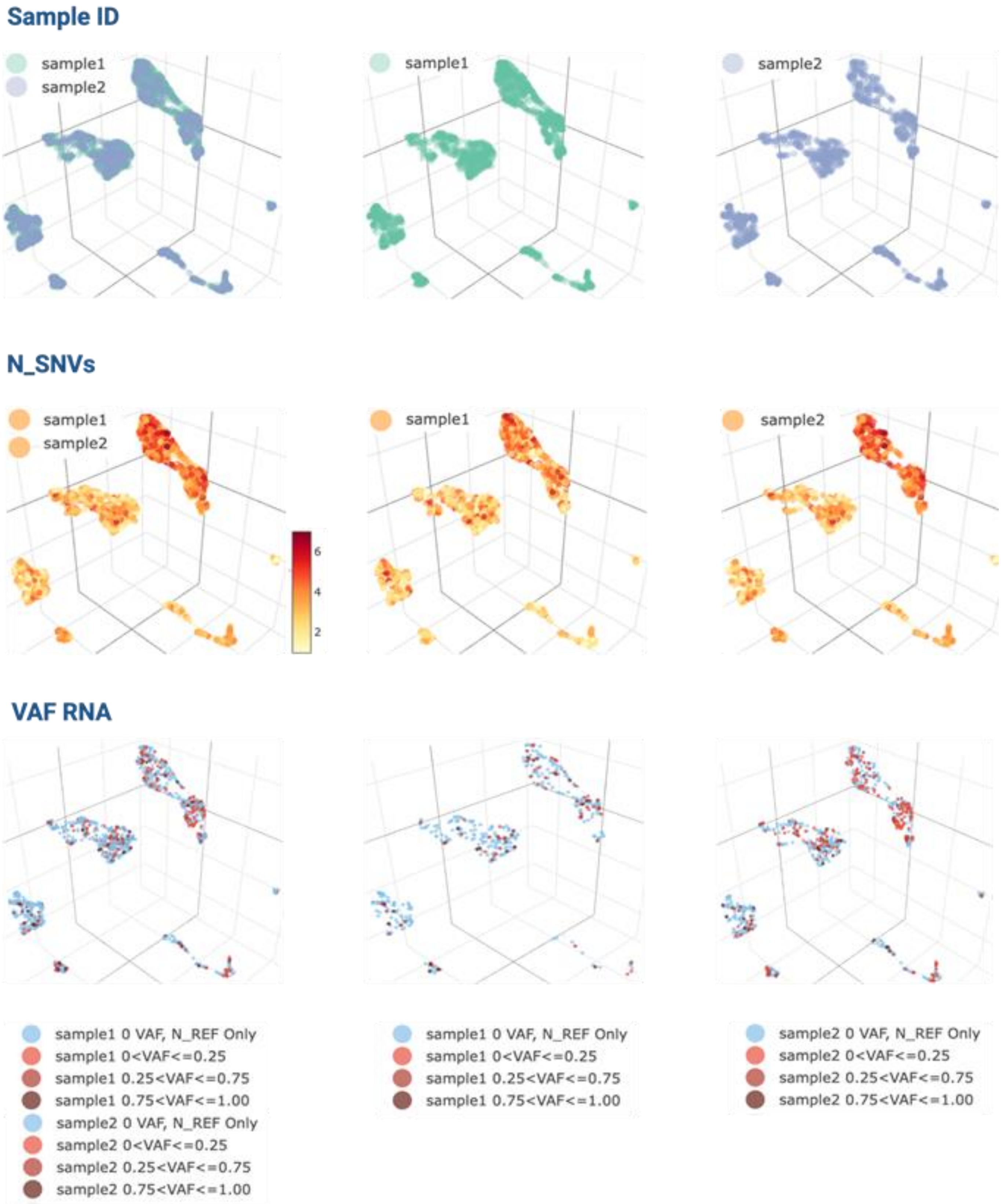


**Supplementary Figure 4**. Visualization of sceSNV metrics from two integrated prostate cancer samples (SAMN16086029 and SAMN16086030) projected into a shared low-dimensional space using Seurat's integration framework via the "enable_integrated" option in scSNViz. The top panel displays a UMAP projection of all cells, allowing selection or exclusion of individual samples to explore integrated versus sample-specific patterns. The middle panel shows the per-cell summary metric N_SNVs (total number of expressed sceSNVs per cell), illustrating intercellular variability in variant burden; integrated and individual sample views are customizable by toggling sample IDs. The bottom panel presents expression of an individual sceSNV (chr17:7403042 C>A, TMEM256) using VAF_RNA, with customizable display of integrated or sample-specific patterns via sample ID selection.

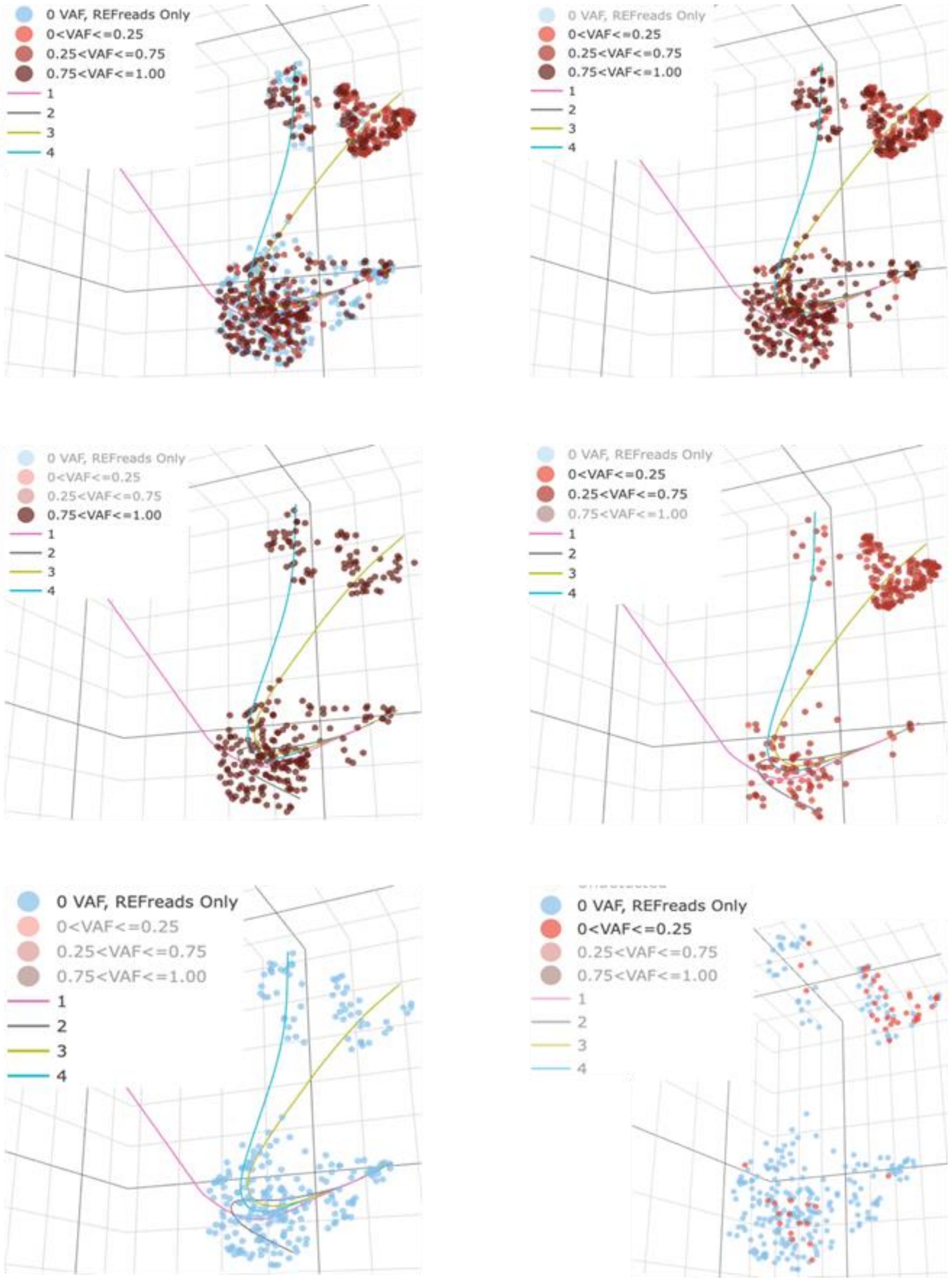


**Supplementary Figure 5**. Interactive visualization capabilities of scSNViz demonstrated in UMAP space using sample SAMN13012147 (cholangiocarcinoma) and a heterozygous germline sceSNV (2:85394936 T>C) in *CAPG* (also shown in Figure 1c). This example demonstrates the breadth of dynamic filtering and visualization options available in **scSNViz**. Panels illustrate: (**top left**) cells expressing the gene harboring the variant, with either the reference, variant allele, or both (cells not expressing the sceSNV-harboring gene are not shown); (**top right)** only cells expressing the variant allele; (**middle left**) cells with near-monoallelic expression of the variant allele; (**middle right**) cells exhibiting low-level variant expression, which appear spatially clustered; (**bottom left**) cells expressing only the reference allele; and (**bottom right**) optional overlay of **Slingshot** trajectories for lineage inference.

**SAMN12799270, neuroblastoma**
**6:26104289_T>C, *H4C3,* 3-prime-UTR**

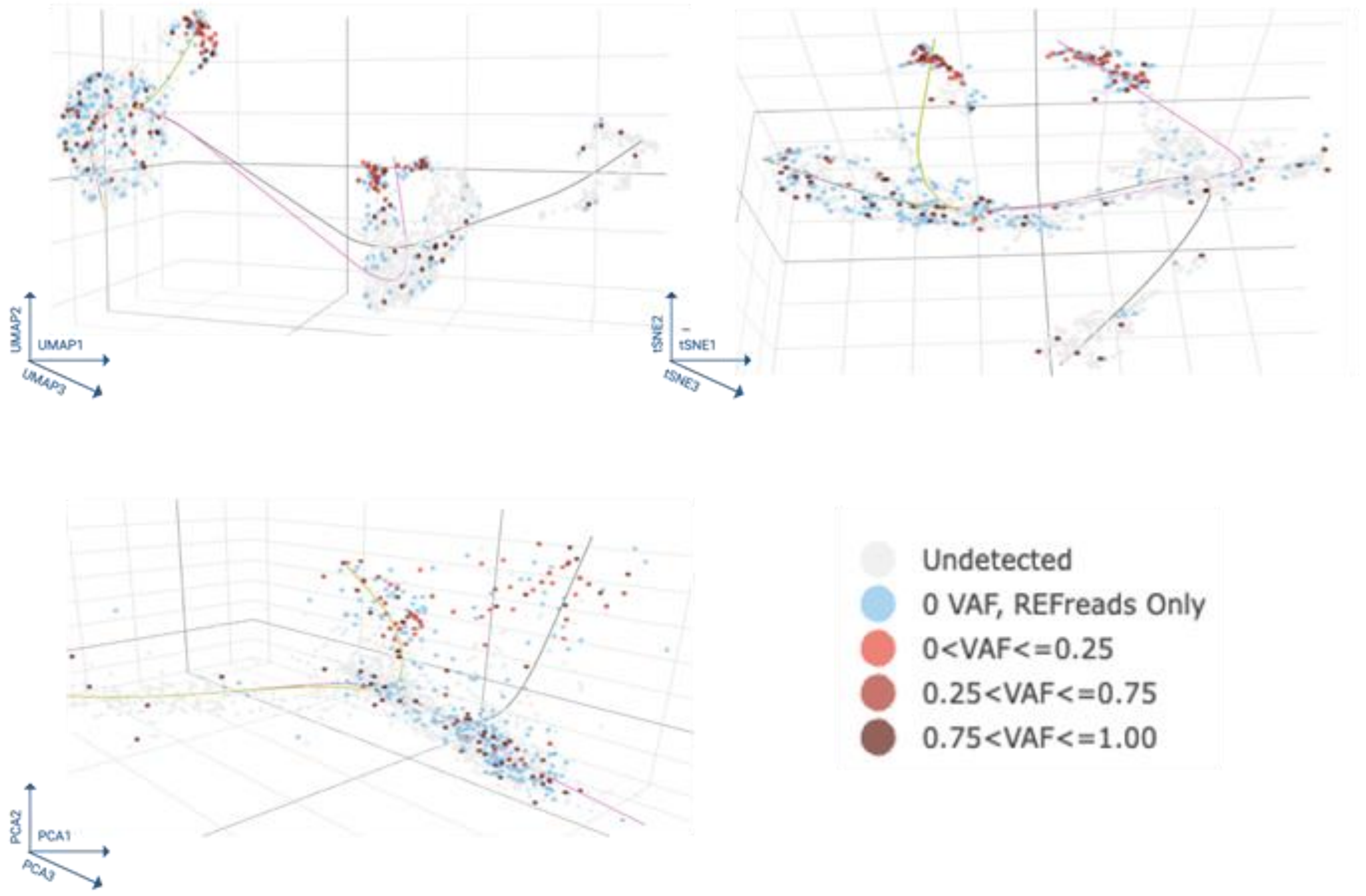


**Supplementary Figure 6 scSNViz** offers visualization on UMAP (top left), tSNE (top right), and PCA (bottom left) dimensionally reduced cell projections.

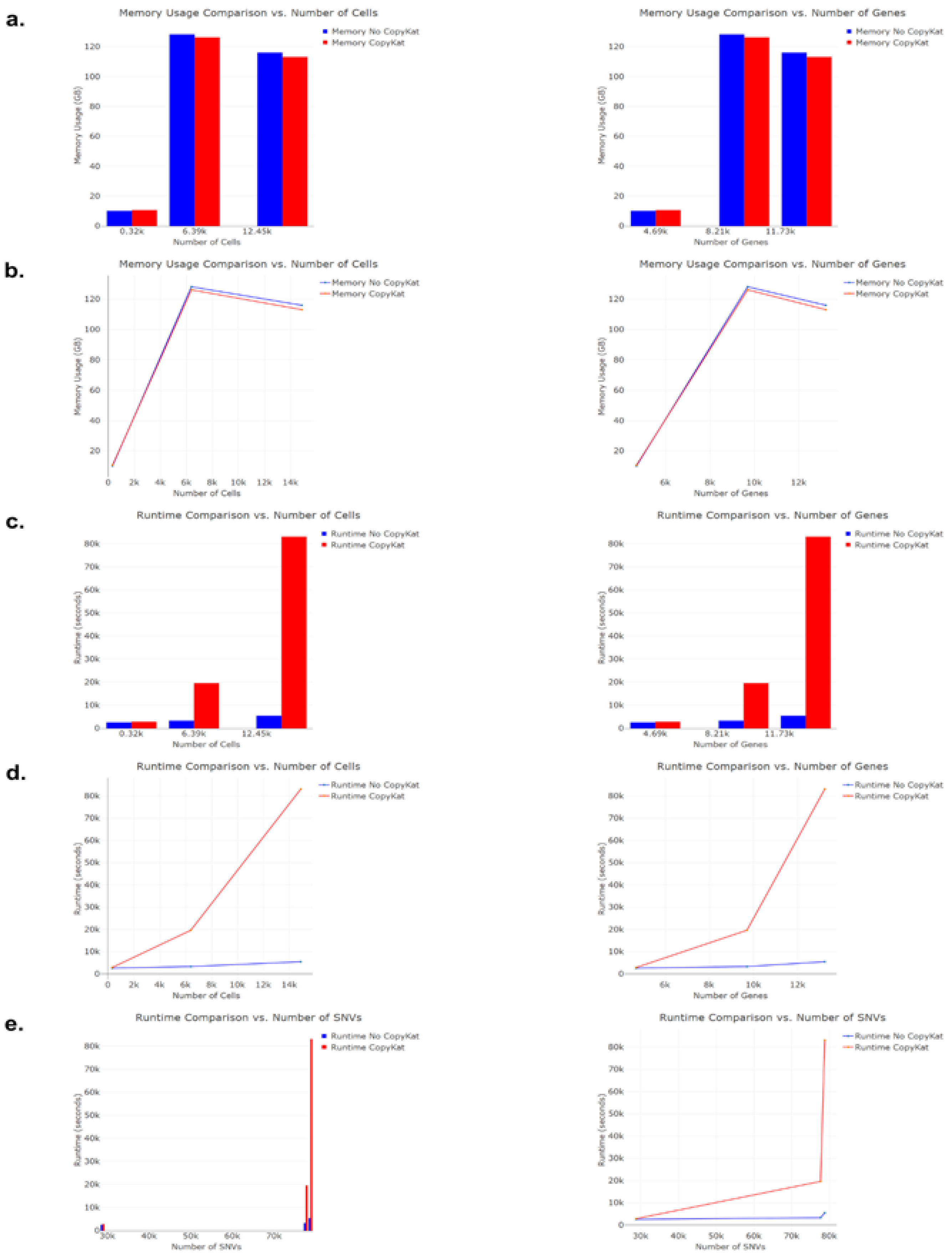


**Supplementary Figure 7 .** Total memory usage was primarily determined by the number of features and cells in the input matrix, as well as the size of the SNV matrix. Runtime was also largely influenced by the number of cells and features, with a notable increase associated with the use of **CopyKat**, ranging from an additional 5 minutes to up to 20 hours. This overhead can be mitigated by enabling **parallel processing** within the CopyKat tool - an option not utilized in this analysis but recommended for large-scale datasets.

COSV63506293, SAMN13822233, nsclc

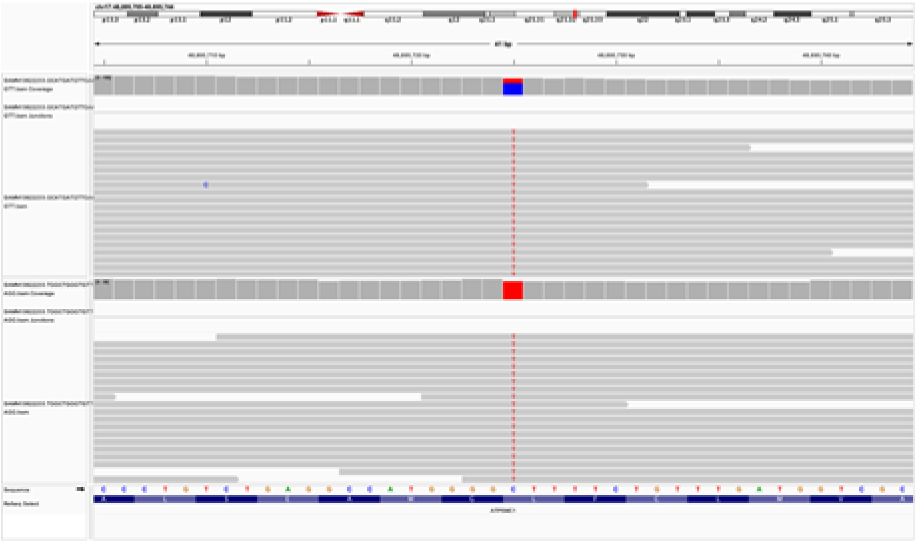

COSV63506293, SAMN12799259, pc

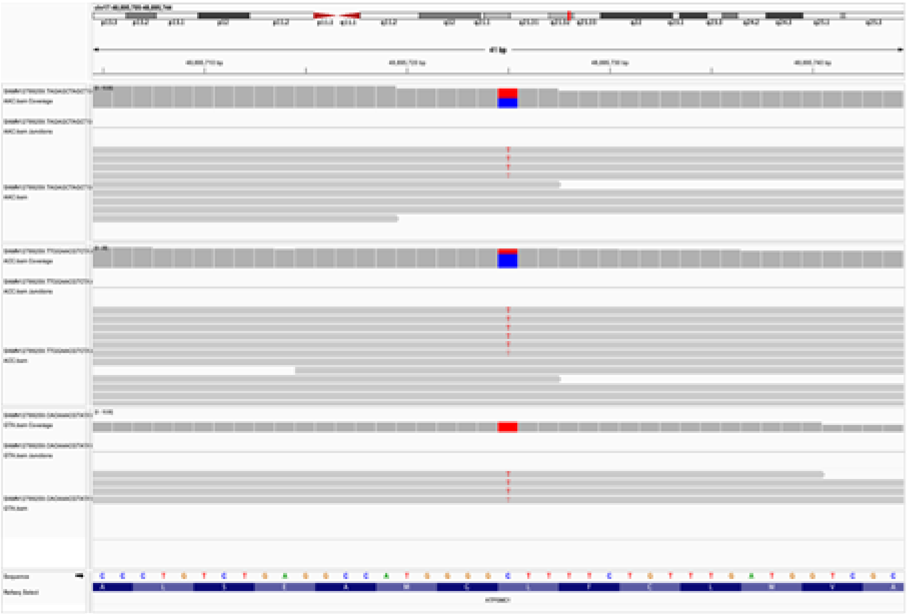

COSV63506293, SAMN13012150, chlg

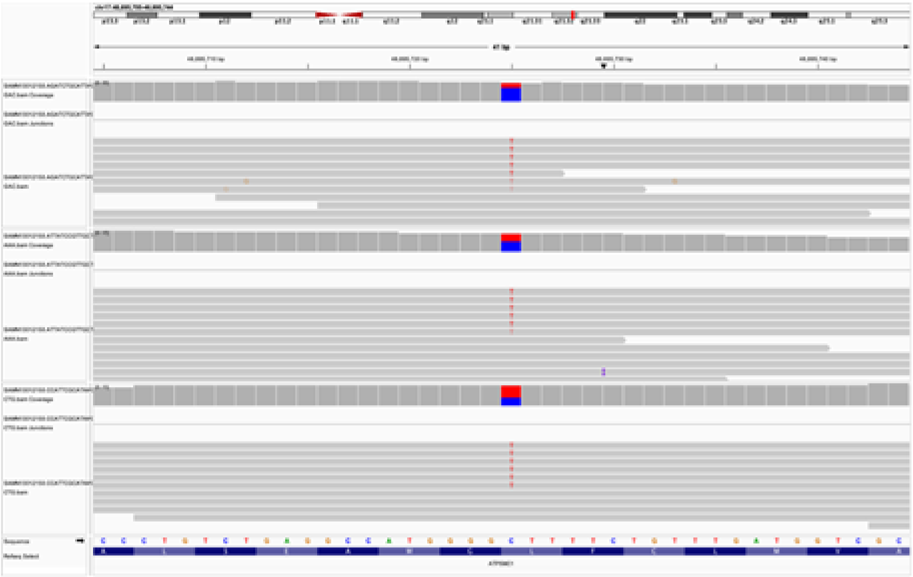

**Supplementary Figure 8.** Integrative Genomics Viewer (IGV) visualization of cell-level scRNA-seq alignments at a genomic locus reported as a somatic mutation in COSMIC (COSV63506293; 17:48895725 C>T in *ATP5MC1*). Across all examined cells and samples, high-quality variant-supporting reads and alignment patterns are observed, consistent with a likely true positive sceSNV.

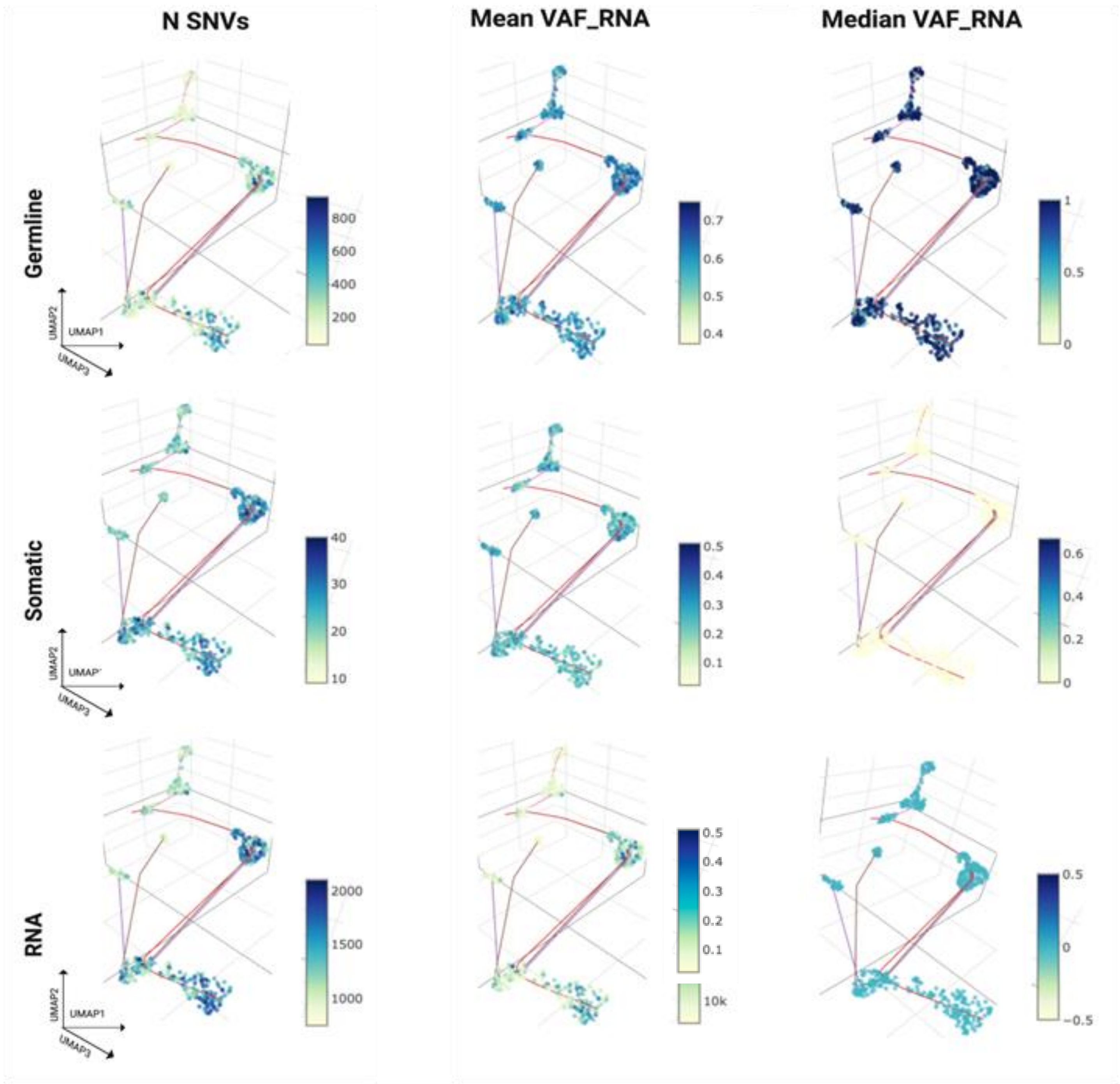


**Supplementary Figure 9** UMAP projections of sceSNVs categorized as germline (top), somatic (middle), and RNA-origin (bottom) in sample SAMN16086829 (prostate cancer). Each row corresponds to one category of sceSNVs, with panels (left to right) showing: N_SNVs (number of expressed sceSNVs per cell), mean VAF_RNA, and median VAF_RNA calculated across all sceSNVs in the respective set. Distinct VAF_RNA profiles are observed across the three categories, with germline, somatic, and RNA-origin sceSNVs exhibiting characteristic differences in both mean and median VAF_RNA distributions.
