## Supplementary Tables for "scSNViz: Visualization and Analysis of Cell-Specific Expressed SNVs"

Supplementary Table 1. Datasets utilized in the analyses performed with scSNViz.

| ## | NCBI IDs | | Sample info | Chromium Version | Read Length | N_cells | Reference |
|---|---|---|---|---|---|---|---|
| | PRJNA | SAMN | tissue | | | | |
| 1 | PRJNA662503 | SAMN16086830 | Prostate cancer | v2 | 150 | 1455 | *Ma et al., 2020* |
| 2 | | SAMN16086829 | | v2 | 150 | 2019 | |
| 3 | PRJNA600483 | SAMN13822232 | Non-small Cell Lung Carcinoma | v2 | 150 | 4074 | *Wang et al., 2019* |
| 4 | | SAMN13822233 | | v2 | 150 | 8381 | |
| 5 | | SAMN13822234 | | v2 | 150 | 7826 | |
| 6 | PRJNA576876 | SAMN13012145 | Intrahepatic Cholangiocarcinoma | v2 | 150 | 3519 | *Zhang et al.,2020* |
| 7 | | SAMN13012146 | | v2 | 150 | 2453 | |
| 8 | | SAMN13012147 | | v2 | 150 | 3579 | |
| 9 | | SAMN13012148 | | v2 | 150 | 3769 | |
| 10 | | SAMN13012149 | | v2 | 150 | 2738 | |
| 11 | | SAMN13012150 | | v2 | 150 | 4993 | |
| 12 | PRJNA573097 | SAMN12799275 | Neuroblastoma | v2 | 150 | 4068 | *Dong et al., 2020* |
| 13 | | SAMN12799274 | | v2 | 150 | 5789 | |
| 14 | | SAMN12799273 | | v2 | 150 | 6988 | |
| 15 | | SAMN12799272 | | v2 | 150 | 2997 | |
| 16 | | SAMN12799270 | | v2 | 150 | 6836 | |
| 17 | | SAMN12799269 | | v2 | 150 | 6994 | |
| 18 | | SAMN12799266 | | v2 | 150 | 12488 | |
| 19 | | SAMN12799264 | | v2 | 150 | 16554 | |
| 20 | | SAMN12799263 | | v2 | 150 | 4273 | |
| 21 | | SAMN15453063 | | v3 | 150 | 12441 | |
| 22 | | SAMN15453064 | | v3 | 150 | 7582 | |
| 23 | PRJNA573097 | SAMN12799261 | Normal Fetal Adrenal | v2 | 150 | 9112 | |
| 24 | | SAMN12799259 | | v2 | 150 | 4916 | |
| 25 | | SAMN12799258 | | v2 | 150 | 26329 | |
| 26 | | SAMN12799257 | | v2 | 150 | 21816 | |
| 27 | PRJNA573097 | SAMN15453062 | Normal Embryo | v3 | 150 | 19639 | |
| 28 | | SAMN15453069 | | v3 | 150 | 14375 | |
| | | | | | **Total** | **228003** | |

**Supplementary Table 2.** Set of sceSNVs used to demonstrate set-level visualization features in scSNViz (Figure 1b)

| ## | SNV | geneList | accession | functionGVS | rsID | Aas | genomesESP | SNV_COS_ID |
|---|---|---|---|---|---|---|---|---|
| 1 | 1:1014228_G>A | ISG15 | NM_005101.4 | missense | 1921 | SER,ASN | A=5223/G=7783 | COSV65106391 |
| 2 | 1:1014274_A>G | ISG15 | NM_005101.4 | synonymous | 8997 | VAL | G=10668/A=2336 | COSV65106735 |
| 3 | 1:109737079_C>T | GSTM3 | NM_000849.5 | missense | 7483 | VAL,ILE | T=3043/C=9963 | COSV56661922 |
| 4 | 1:111243307_C>A | CHI3L2 | NM_001025197.1 | 3-prime-UTR | 8535 | none | unknown | |
| 5 | 1:111478407_A>G | C1orf162 | NM_001300834.2 | 3-prime-UTR | 1054680 | none | unknown | |
| 6 | 1:11750708_T>A | AGTRAP | NM_001040194.1 | 3-prime-UTR | 6540993 | none | unknown | |
| 7 | 1:155087375_A>G | EFNA3 | NM_004952.5 | 3-prime-UTR | 2306124 | none | unknown | |
| 8 | 1:161213726_C>T | NDUFS2 | NM_001166159.2 | synonymous | 1136207 | ALA | T=1378/C=11628 | COSV57154020 |
| 9 | 1:167430725_T>C | CD247 | NM_000734.4 | 3-prime-UTR | 947480 | none | unknown | |
| 10 | 1:167430837_T>A | CD247 | NM_000734.4 | 3-prime-UTR | 1052231 | none | unknown | |
| 11 | 1:168055440_G>A | DCAF6 | NM_001017977.2 | intron | 16865519 | none | unknown | |
| 12 | 1:192812042_C>G | RGS2 | NM_002923.4 | 3-prime-UTR | 4606 | none | unknown | |
| 13 | 1:202005973_G>A | RNPEP,ELF3-AS1 | NM_001319182.2 | 3-prime-UTR | 117567081 | none | unknown | |
| 14 | 1:212445997_T>C | NENF | NM_013349.5 | synonymous | 4804 | ASP | C=5623/T=7383 | COSV65340955 |
| 15 | 1:22661465_A>G | C1QB | NM_000491.5 | 3-prime-UTR | 10580 | none | unknown | |
| 16 | 1:228494629_C>G | RNF187 | NM_001010858.3 | 3-prime-UTR | 0 | none | unknown | |
| 17 | 1:25900804_G>A | STMN1 | NM_001145454.3 | intron | 0 | none | unknown | |
| 18 | 1:77773018_A>G | none | none | intergenic | 0 | none | unknown | |
| 19 | 1:77773045_T>C | none | none | intergenic | 0 | none | unknown | |
| 20 | 1:77773058_A>G | none | none | intergenic | 0 | none | unknown | |
| 21 | 1:944307_T>C | NOC2L,SAMD11 | NM_015658.4 | 3-prime-UTR | 2839 | none | unknown | COSV58988633 |
| 22 | 10:133362587_G>A | ECHS1 | NM_004092.4 | 3-prime-UTR | 4604 | none | unknown | |
| 23 | 10:17237563_G>T | VIM | NM_003380.5 | 3-prime-UTR | 1049341 | none | unknown | |
| 24 | 10:3136580_C>G | PFKP | NM_001242339.1 | 3-prime-UTR | 184205778 | none | C=13006 | |
| 25 | 10:3136623_G>C | PFKP | NM_001242339.1 | 3-prime-UTR | 9063 | none | C=1569/G=11437 | COSV56538751 |
| 26 | 10:3136716_T>C | PFKP | NM_001242339.1 | 3-prime-UTR | 542 | none | unknown | COSV56525277 |
| 27 | 10:68342992_C>A | HNRNPH3,RUFY2 | NM_001322434.1 | 3-prime-UTR | 3199937 | none | unknown | |
| 28 | 11:16755795_G>A | C11orf58 | NM_014267.6 | 3-prime-UTR | 4576801 | none | unknown | |
| 29 | 11:1892147_A>G | LSP1 | NM_001013253.2 | 3-prime-UTR | 548195 | none | unknown | |
| 30 | 11:46383776_C>A | MDK | NM_001012333.2 | 3-prime-UTR | 116869512 | none | unknown | COSV58994791 |
| 31 | 11:47354897_G>A | SPI1 | NM_001080547.2 | 3-prime-UTR | 1057233 | none | unknown | |
| 32 | 11:67585218_A>G | GSTP1 | NM_000852.4 | missense | 1695 | ILE,VAL | G=4535/A=8033 | COSV66992376 |
| 33 | 11:78079607_T>C | NDUFC2,NDUFC2-KCTD14 | NM_001203260.2 | synonymous | 534418 | LEU | C=11218/T=1602 | COSV55247502 |
| 34 | 11:93536126_G>A | SMCO4 | NM_020179.3 | intron | 0 | none | unknown | |
| 35 | 12:132704876_G>A | PXMP2 | NM_018663.3 | 3-prime-UTR | 10007 | none | unknown | |
| 36 | 12:55725474_C>T | CD63 | NM_001257389.1 | 3-prime-UTR | 1037113923 | none | unknown | COSV57677305 |
| 37 | 12:55725822_A>G | CD63 | NM_001257389.1 | synonymous | 0 | ALA | A=13006 | |
| 38 | 12:55757507_T>C | SARNP | NM_033082.4 | 3-prime-UTR | 7068 | none | C=2854/T=10152 | COSV60242616 |
| 39 | 12:9598073_A>G | KLRB1 | NM_002258.3 | missense | 1135816 | ILE,THR | G=4425/A=8565 | COSV57590625 |
| 40 | 14:103519918_C>T | CKB | NM_001362531.2 | synonymous | 1803283 | GLU | T=7592/C=5410 | COSV62379683 |
| 41 | 14:105488785_G>A | CRIP1 | NM_001311.5 | 3-prime-UTR | 0 | none | unknown | |
| 42 | 14:20782207_T>C | RNASE6 | NM_005615.5 | 3-prime-UTR | 7156801 | none | unknown | |
| 43 | 14:35402011_G>A | NFKBIA | NM_020529.3 | 3-prime-UTR | 8904 | none | A=5786/G=7220 | COSV53754079 |
| 44 | 14:49586377_G>A | RPS29 | NM_001030001.4 | 5-prime-UTR | 1282480270 | none | G=13006 | COSV99817971 |
| 45 | 14:77708130_A>C | SLIRP | NM_001267863.1 | synonymous | 11159286 | ARG | C=12016/A=990 | COSV53174819 |
| 46 | 14:98972797_C>T | none | none | intergenic | 372490154 | none | unknown | |
| 47 | 15:34341832_A>G | NOP10 | NM_018648.4 | 3-prime-UTR | 3063 | none | unknown | COSV60995282 |
| 48 | 15:34341923_C>G | NOP10 | NM_018648.4 | 3-prime-UTR | 1045238 | none | G=2052/C=10946 | COSV60995138 |
| 49 | 15:34341937_G>A | NOP10 | NM_018648.4 | 3-prime-UTR | 1045204 | none | A=1736/G=11262 | COSV60995143 |
| 50 | 15:34341938_T>C | NOP10 | NM_018648.4 | 3-prime-UTR | 1045194 | none | C=1928/T=11070 | COSV60995151 |

Supplementary Table 2 (cont).

| ## | SNV | geneList | accession | functionGVS | rsID | Aas | genomesESP | SNV_COS_ID |
|---|---|---|---|---|---|---|---|---|
| 51 | 15:44717976_G>A | B2M | NM_004048.3 | 3-prime-UTR | 0 | none | unknown | |
| 52 | 15:48878780_G>A | EID1,SHC4 | NM_014335.3 | 3-prime-UTR | 16961791 | none | unknown | |
| 53 | 15:69452821_C>T | RPLP1 | NM_001003.3 | 5-prime-UTR | 529326572 | none | unknown | |
| 54 | 15:81308981_A>G | IL16 | NM_001172128.2 | 3-prime-UTR | 859 | none | unknown | |
| 55 | 15:88652295_A>G | ISG20 | NM_001303233.2 | synonymous | 1137166 | LEU | G=10738/A=2260 | COSV60143565 |
| 56 | 15:92898037_G>A | LINC01578 | NR_037600.1 | non-coding-exo | 7743 | none | unknown | |
| 57 | 16:1325262_T>G | UBE2I | NM_003345.5 | 3-prime-UTR | 7302 | none | T=4566 | |
| 58 | 16:57275_T>G | SNRNP25 | NM_024571.4 | 3-prime-UTR | 1045001 | none | unknown | COSV51954838 |
| 59 | 16:84565600_C>T | COTL1 | NM_021149.5 | 3-prime-UTR | 0 | none | unknown | |
| 60 | 16:84565720_G>A | COTL1 | NM_021149.5 | 3-prime-UTR | 774809134 | none | unknown | |
| 61 | 16:85922438_A>T | IRF8 | NM_001363907.1 | 3-prime-UTR | 6638 | none | unknown | |
| 62 | 16:87483044_A>T | ZCCHC14 | NM_015144.3 | intron | 11864816 | none | unknown | |
| 63 | 17:15229806_C>A | PMP22 | NM_000304.4 | 3-prime-UTR | 7415 | none | unknown | COSV56601929 |
| 64 | 17:5499637_A>G | LOC728392 | NM_001162371.3 | 3-prime-UTR | 5862 | none | unknown | |
| 65 | 17:68532637_C>G | PRKAR1A,FAM20A | NM_001276289.1 | 3-prime-UTR | 6958 | none | unknown | COSV56837023 |
| 66 | 17:7014384_C>T | RNASEK,RNASEK-C17orf49 | NM_001004333.4 | 3-prime-UTR | 7338 | none | unknown | COSV52351111 |
| 67 | 17:7014479_T>C | RNASEK,RNASEK-C17orf49 | NM_001004333.4 | 3-prime-UTR | 12135 | none | unknown | |
| 68 | 17:7240720_G>A | GABARAP | NM_007278.2 | 3-prime-UTR | 0 | none | unknown | |
| 69 | 18:59158639_T>C | SEC11C | NM_001307941.2 | missense | 0 | PHE,LEU | T=13006 | |
| 70 | 19:10110966_A>G | PPAN,PPAN-P2RY11 | NM_001040664.3 | missense | 11559188 | GLN,ARG | G=613/A=12391 | |
| 71 | 19:10115101_A>G | P2RY11,EIF3G,PPAN-P2RY11 | NM_001040664.3 | 3-prime-UTR | 7401 | none | G=4884/A=8122 | COSV53452898 |
| 72 | 19:1038894_C>T | CNN2 | NM_001303499.2 | 3-prime-UTR | 1057895 | none | unknown | |
| 73 | 19:1106616_T>C | GPX4 | NM_001039847.3 | synonymous | 713041 | LEU | C=7154/T=4930 | COSV62321056 |
| 74 | 19:11450807_A>G | PRKCSH | NM_001001329.2 | 3-prime-UTR | 1056893467 | none | unknown | |
| 75 | 19:11553206_G>A | ELOF1 | NM_001363673.1 | 3-prime-UTR | 72620552 | none | unknown | COSV52947359 |
| 76 | 19:16133516_G>C | RAB8A | NM_005370.5 | 3-prime-UTR | 1043452 | none | unknown | |
| 77 | 19:17320106_G>A | DDA1 | NM_024050.6 | 3-prime-UTR | 10259 | none | unknown | |
| 78 | 19:17320166_G>A | DDA1 | NM_024050.6 | 3-prime-UTR | 1059767 | none | unknown | |
| 79 | 19:17403117_G>C | BST2 | NM_004335.4 | 3-prime-UTR | 13485 | none | unknown | |
| 80 | 19:17788341_G>A | FCHO1 | NM_001161357.2 | 3-prime-UTR | 369047298 | none | A=1/G=13003 | |
| 81 | 19:19201687_C>G | RFXANK,NR2C2AP | NM_001278727.1 | missense | 1802498 | GLN,GLU | G=14/C=12992 | COSV57393414 |
| 82 | 19:2732743_G>A | SLC39A3 | NM_144564.5 | 3-prime-UTR | 9160 | none | A=1254/G=11714 | |
| 83 | 19:2754794_G>A | SGTA | NM_003021.4 | 3-prime-UTR | 7009 | none | unknown | |
| 84 | 19:2754812_C>T | SGTA | NM_003021.4 | 3-prime-UTR | 13282 | none | unknown | |
| 85 | 19:2754980_T>C | SGTA | NM_003021.4 | 3-prime-UTR | 7008 | none | unknown | |
| 86 | 19:33387291_G>A | PEPD | NM_000285.4 | 3-prime-UTR | 77690463 | none | unknown | COSV52287665 |
| 87 | 19:37738477_G>A | ZNF573 | NM_001172689.1 | 3-prime-UTR | 1291 | none | A=8975/G=4031 | COSV59828169 |
| 88 | 19:38878729_T>G | SIRT2 | NM_001193286.1 | 3-prime-UTR | 2015 | none | unknown | |
| 89 | 19:38878874_G>A | SIRT2 | NM_001193286.1 | 3-prime-UTR | 2241703 | none | unknown | |
| 90 | 19:40765199_T>C | SNRPA | NM_004596.5 | 3-prime-UTR | 13108 | none | C=1532/T=11474 | |
| 91 | 19:40796801_C>T | RAB4B,MIA-RAB4B,RAB4B-EG | NM_016154.5 | 3-prime-UTR | 7937 | none | unknown | COSV58279921 |
| 92 | 19:4174401_C>G | SIRT6 | NM_001193285.3 | 3-prime-UTR | 350846 | none | unknown | |
| 93 | 19:44903416_G>A | TOMM40 | NM_001128916.1 | 3-prime-UTR | 10119 | none | unknown | |
| 94 | 19:45475117_A>G | FOSB | NM_001114171.2 | 3-prime-UTR | 1049739 | none | unknown | |
| 95 | 19:45526896_G>A | VASP | NM_003370.4 | 3-prime-UTR | 10995 | none | unknown | |
| 96 | 19:48966669_G>A | FTL | NM_000146.4 | synonymous | 0 | ARG | G=13006 | |
| 97 | 19:48966687_T>G | FTL | NM_000146.4 | synonymous | 0 | ALA | T=13006 | |
| 98 | 19:48966709_G>A | FTL | NM_000146.4 | missense | 768204975 | GLU,LYS | G=13006 | COSV53169512 |
| 99 | 19:48966747_G>C | FTL | NM_000146.4 | 3-prime-UTR | 1432695180 | none | G=13006 | |
| 100 | 19:48966766_G>C | FTL | NM_000146.4 | 3-prime-UTR | 764013805 | none | G=13006 | |

Supplementary Table 2 (cont.)

| ## | SNV | geneList | accession | functionGVS | rsID | Aas | genomesESP | SNV_COS_ID |
|---|---|---|---|---|---|---|---|---|
| 101 | 19:48966787_C>T | FTL | NM_000146.4 | 3-prime-UTR | 771233352 | none | C=7264 | |
| 102 | 19:49447041_C>T | PIH1D1 | NM_017916.3 | missense | 13394 | VAL,ILE | T=10532/C=2474 | COSV51807272 |
| 103 | 19:50797973_G>A | C19orf48 | NM_001290149.1 | 3-prime-UTR | 9991 | none | unknown | COSV54517312 |
| 104 | 19:52734417_T>G | ZNF611 | NM_001161499.2 | intron | 707303 | none | unknown | |
| 105 | 19:55643352_G>A | ZNF580,ZNF581 | NM_001163423.1 | 3-prime-UTR | 11233 | none | unknown | COSV54402651 |
| 106 | 19:58394796_T>C | RPS5 | NM_001009.4 | 3-prime-UTR | 2241787 | none | C=5824/T=7182 | COSV52190570 |
| 107 | 19:58582097_C>T | none | none | intergenic | 0 | none | unknown | |
| 108 | 19:58582117_G>T | none | none | intergenic | 0 | none | unknown | |
| 109 | 19:7116272_G>C | INSR | NM_000208.4 | 3-prime-UTR | 1051651 | none | unknown | |
| 110 | 19:7647391_A>G | STXBP2 | NM_001127396.3 | missense | 6791 | ILE,VAL | G=8655/A=4315 | COSV55401668 |
| 111 | 19:8311547_G>A | NDUFA7 | NM_005001.5 | synonymous | 561 | PRO | A=1819/G=10649 | COSV56846606 |
| 112 | 19:896740_C>G | R3HDM4 | NM_138774.4 | 3-prime-UTR | 7969 | none | unknown | COSV54122929 |
| 113 | 2:112829856_C>G | IL1B | NM_000576.3 | 3-prime-UTR | 1071676 | none | unknown | |
| 114 | 2:119372544_T>A | DBI | NM_001079862.3 | 3-prime-UTR | 0 | none | unknown | |
| 115 | 2:174720839_T>A | none | none | intergenic | 0 | none | unknown | |
| 116 | 2:174720854_G>A | none | none | intergenic | 0 | none | unknown | |
| 117 | 2:232480018_G>C | ECEL1 | NM_001290787.2 | 3-prime-UTR | 2741278 | none | unknown | COSV58811853 |
| 118 | 2:241495187_A>G | STK25 | NM_001271977.2 | 3-prime-UTR | 7618 | none | unknown | |
| 119 | 2:241676047_T>C | DTYMK | NM_001165031.2 | 3-prime-UTR | 5860 | none | unknown | COSV59870962 |
| 120 | 2:26190948_C>T | HADHA,GAREM2 | NM_000182.5 | 3-prime-UTR | 1049987 | none | unknown | |
| 121 | 2:277995_A>G | ACP1 | NM_004300.4 | 3-prime-UTR | 6855 | none | unknown | |
| 122 | 2:47160272_A>G | CALM2 | NM_001305624.1 | 3-prime-UTR | 0 | none | unknown | |
| 123 | 2:47160299_C>T | CALM2 | NM_001305624.1 | 3-prime-UTR | 11551467 | none | unknown | |
| 124 | 2:85581901_G>A | VAMP8 | NM_003761.5 | 3-prime-UTR | 0 | none | unknown | |
| 125 | 2:85698725_C>G | GNLY | NM_001302758.2 | 3-prime-UTR | 12845 | none | unknown | COSV55702702 |
| 126 | 20:18763739_T>C | DTD1 | NM_080820.6 | 3-prime-UTR | 9139 | none | unknown | |
| 127 | 20:24959885_A>G | CST7 | NM_003650.4 | 3-prime-UTR | 1056036 | none | unknown | |
| 128 | 20:4024544_C>T | none | none | intergenic | 878976490 | none | unknown | |
| 129 | 20:4024575_G>A | none | none | intergenic | 6084555 | none | unknown | |
| 130 | 20:4024581_C>T | none | none | intergenic | 8122653 | none | unknown | |
| 131 | 20:46016514_T>C | MMP9,SLC12A5-AS1 | NM_004994.3 | 3-prime-UTR | 9509 | none | unknown | |
| 132 | 20:5925133_C>A | CHGB | NM_001819.3 | 3-prime-UTR | 2821 | none | unknown | |
| 133 | 20:62387103_C>T | RPS21 | NM_001024.4 | 5-prime-UTR | 972475308 | none | unknown | |
| 134 | 21:44886223_G>T | ITGB2 | NM_000211.5 | 3-prime-UTR | 1160263 | none | unknown | COSV56609771 |
| 135 | 22:19435833_G>T | MRPL40 | NM_001318151.2 | synonymous | 77131689 | ARG | T=2/G=13004 | |
| 136 | 22:32532118_T>C | SYN3 | NM_001135774.2 | intron | 10460755 | none | unknown | |
| 137 | 22:41663764_G>T | XRCC6 | NM_001288976.2 | synonymous | 132788 | GLY | T=3487/G=9519 | COSV63752372 |
| 138 | 22:43162920_A>G | TSPO | NM_000714.6 | missense | 6971 | THR,ALA | G=9431/A=3553 | COSV53355454 |
| 139 | 22:43163131_G>T | TSPO | NM_000714.6 | 3-prime-UTR | 6973 | none | unknown | |
| 140 | 22:49963768_C>T | PIM3 | NM_001001852.4 | 3-prime-UTR | 13811 | none | unknown | |
| 141 | 22:50525807_G>A | TYMP,SCO2 | NM_001113755.3 | missense | 11479 | SER,LEU | A=846/G=11530 | COSV52688025 |
| 142 | 22:50526017_A>T | TYMP,SCO2 | NM_001113755.3 | synonymous | 1138404 | GLY | unknown | |
| 143 | 3:129314956_C>A | H1-10 | NM_006026.4 | 3-prime-UTR | 0 | none | unknown | |
| 144 | 3:169966810_C>G | SEC62 | NM_003262.4 | 5-prime-UTR | 375131407 | none | T=1/C=12563 | |
| 145 | 3:46922012_T>C | CCDC12 | NM_001277074.1 | 3-prime-UTR | 4587 | none | C=9269/T=3737 | COSV52765225 |
| 146 | 4:108625210_T>A | RPL34 | NM_000995.4 | stop-lost | 0 | stop,LYS | T=12978 | |
| 147 | 4:17486621_G>A | QDPR | NM_000320.3 | 3-prime-UTR | 1031327 | none | unknown | |
| 148 | 4:17486663_T>G | QDPR | NM_000320.3 | 3-prime-UTR | 699460 | none | unknown | |
| 149 | 4:17486723_G>A | QDPR | NM_000320.3 | 3-prime-UTR | 1049601 | none | unknown | |
| 150 | 4:17486728_T>C | QDPR | NM_000320.3 | 3-prime-UTR | 1049600 | none | unknown | |

### Supplementary Table 2 (cont.)

| ## | SNV | geneList | accession | functionGVS | rsID | Aas | genomesESP | SNV_COS_ID |
|---|---|---|---|---|---|---|---|---|
| 151 | 4:6642664_G>A | MRFAP1 | NM_001272053.1 | 3-prime-UTR | 1059220 | none | unknown | |
| 152 | 4:87982701_C>T | SPP1 | NM_000582.2 | synonymous | 1126616 | ALA | T=3124/C=9882 | COSV52951697 |
| 153 | 4:87983190_A>C | SPP1 | NM_000582.2 | 3-prime-UTR | 9138 | none | unknown | |
| 154 | 5:10264964_C>T | CCT5 | NM_001306153.1 | 3-prime-UTR | 699113 | none | unknown | |
| 155 | 5:177306115_A>G | PRELID1,MXD3 | NM_001142935.2 | 3-prime-UTR | 4631 | none | G=11724/A=1282 | COSV57462840 |
| 156 | 5:177306615_T>C | PRELID1,MXD3 | NM_001142935.2 | 3-prime-UTR | 9834 | none | C=11645/T=1173 | COSV57463085 |
| 157 | 5:35871088_G>A | IL7R | NM_002185.5 | missense | 1494555 | VAL,ILE | A=9700/G=3306 | COSV57406117 |
| 158 | 5:53683267_G>A | NDUFS4 | NM_001318051.2 | 3-prime-UTR | 567 | none | A=5456/G=7550 | COSV57019229 |
| 159 | 6:111764035_A>G | FYN | NM_002037.5 | intron | 6933144 | none | unknown | COSV57611981 |
| 160 | 6:122725777_A>G | PKIB | NM_001270393.1 | 3-prime-UTR | 1132635 | none | unknown | COSV57794791 |
| 161 | 6:132814568_C>G | none | none | upstream-gene | 576858600 | none | unknown | |
| 162 | 6:132814747_A>G | RPS12 | NM_001016.4 | 5-prime-UTR | 9483504 | none | G=6929/A=6077 | COSV57767099 |
| 163 | 6:149811183_A>T | PCMT1 | NM_001252049.1 | 3-prime-UTR | 4552 | none | unknown | COSV66308362 |
| 164 | 6:159778994_A>C | ACAT2,TCP1 | NM_001008897.1 | 3-prime-UTR | 4832 | none | C=2316/A=10682 | COSV58458988 |
| 165 | 6:26104052_T>C | H4C3 | NM_003542.4 | synonymous | 2229768 | ILE | C=2952/T=10054 | COSV58090568 |
| 166 | 6:26104289_T>C | H4C3 | NM_003542.4 | 3-prime-UTR | 0 | none | T=13006 | |
| 167 | 6:26104297_C>A | H4C3 | NM_003542.4 | 3-prime-UTR | 1280403614 | none | C=13006 | |
| 168 | 6:26104303_C>A | H4C3 | NM_003542.4 | 3-prime-UTR | 1205348791 | none | C=13006 | |
| 169 | 6:29555869_G>C | UBD | NM_006398.4 | 3-prime-UTR | 444013 | none | C=5024/G=3416 | |
| 170 | 6:29555899_C>G | UBD | NM_006398.4 | missense | 8337 | CYS,SER | G=6756/C=1682 | |
| 171 | 6:31268757_G>A | HLA-C | NM_002117.6 | 3-prime-UTR | 35075694 | none | unknown | |
| 172 | 6:31268790_T>C | HLA-C | NM_002117.6 | 3-prime-UTR | 1049281 | none | unknown | |
| 173 | 6:31268866_T>C | HLA-C | NM_002117.6 | 3-prime-UTR | 1130552 | none | unknown | |
| 174 | 6:31268902_T>C | HLA-C | NM_002117.6 | 3-prime-UTR | 1130580 | none | unknown | COSV66115208 |
| 175 | 6:31268913_T>G | HLA-C | NM_002117.6 | 3-prime-UTR | 1130592 | none | unknown | COSV66119531 |
| 176 | 6:31616832_G>A | AIF1 | NM_001318970.2 | missense | 753881781 | GLU,LYS | G=13006 | |
| 177 | 6:31616909_G>A | AIF1 | NM_001318970.2 | 3-prime-UTR | 0 | none | G=13006 | |
| 178 | 6:31669957_A>T | CSNK2B | NM_001282385.1 | 3-prime-UTR | 5872 | none | T=1883/A=6557 | COSV65507945 |
| 179 | 6:31730686_C>T | CLIC1 | NM_001287593.1 | 3-prime-UTR | 0 | none | unknown | |
| 180 | 6:36602589_T>C | SRSF3 | NM_003017.5 | 3-prime-UTR | 7344 | none | unknown | COSV59672440 |
| 181 | 6:37483138_T>C | CCDC167 | NM_138493.3 | 3-prime-UTR | 10692 | none | C=9630/T=3376 | COSV64982259 |
| 182 | 6:85678170_G>T | SNHG5 | NR_003038.2 | non-coding-exc | 1059307 | none | unknown | |
| 183 | 6:85678734_A>C | none | none | upstream-gene | 1207944590 | none | unknown | |
| 184 | 7:150338437_T>G | RARRES2 | NM_002889.4 | 3-prime-UTR | 4721 | none | unknown | COSV56231044 |
| 185 | 7:150791528_A>G | TMEM176B | NM_001101311.1 | 3-prime-UTR | 2302479 | none | G=8453/A=4553 | COSV58439096 |
| 186 | 7:24285390_G>A | NPY | NM_000905.4 | synonymous | 5573 | SER | A=5945/G=7061 | COSV54215135 |
| 187 | 7:42937596_G>A | MRPL32 | NM_031903.3 | 3-prime-UTR | 631561 | none | A=12789/G=213 | |
| 188 | 8:123015388_T>C | DERL1 | NM_001134671.2 | 3-prime-UTR | 7159 | none | unknown | COSV52364774 |
| 189 | 8:22594175_G>C | PDLIM2 | NM_001368120.1 | 3-prime-UTR | 3064 | none | unknown | COSV56133534 |
| 190 | 8:32094528_A>T | NRG1,NRG1-IT1 | NM_001159995.3 | intron | 16878794 | none | unknown | |
| 191 | 8:79666159_T>A | STMN2 | XM_005251142.2 | 3-prime-UTR | 904037857 | none | unknown | |
| 192 | 9:127451406_G>C | RPL12 | NM_000976.4 | 5-prime-UTR | 747333236 | none | unknown | |
| 193 | 9:137273425_T>G | NELFB | NM_015456.5 | 3-prime-UTR | 8281 | none | unknown | |
| 194 | 9:36211830_C>T | CLTA | NM_001076677.3 | 3-prime-UTR | 1053414 | none | unknown | COSV54254584 |
| 195 | 9:93121503_T>C | NINJ1 | NM_004148.4 | 3-prime-UTR | 7033638 | none | unknown | |
| 196 | 9:93121564_G>A | NINJ1 | NM_004148.4 | 3-prime-UTR | 12238760 | none | unknown | |
| 197 | 9:93121676_A>T | NINJ1 | NM_004148.4 | 3-prime-UTR | 1127851 | none | unknown | |
| 198 | X:23786051_C>T | SAT1 | NM_002970.3 | 3-prime-UTR | 0 | none | unknown | |
| 199 | X:23786107_G>A | SAT1 | NM_002970.3 | 3-prime-UTR | 0 | none | unknown | |
| 200 | X:47585586_T>C | SYN1,TIMP1 | NM_003254.3 | synonymous | 4898 | PHE | C=4892/T=5671 | COSV54483037 |
| 201 | X:81298369_A>T | SH3BGRL | NM_003022.3 | 3-prime-UTR | 149428539 | none | unknown | |
